# High refresh rate display for natural monocular viewing in AOSLO psychophysics experiments

**DOI:** 10.1101/2024.05.26.595808

**Authors:** Benjamin Moon, Glory Linebach, Angelina Yang, Samantha K. Jenks, Michele Rucci, Martina Poletti, Jannick P. Rolland

**Affiliations:** The Institute of Optics, University of Rochester, Rochester, NY 14627, USA; Center for Visual Science, University of Rochester, Rochester, NY 14627, USA; Department of Brain and Cognitive Sciences, University of Rochester, Rochester, NY 14627, USA; Department of Neuroscience, University of Rochester, Rochester, NY 14627, USA; Department of Biomedical Engineering, University of Rochester, Rochester, NY 14627, USA

## Abstract

By combining an external display operating at 360 frames per second with an Adaptive Optics Scanning Laser Ophthalmoscope (AOSLO) for human foveal imaging, we demonstrate color stimulus delivery at high spatial and temporal resolution in AOSLO psychophysics experiments. A custom pupil relay enables viewing of the stimulus through a 3-mm effective pupil diameter and provides refractive error correction from -8 to +4 diopters. Performance of the assembled and aligned pupil relay was validated by measuring the wavefront error across the field of view and correction range, and the as-built Strehl ratio was 0.64 or better. High-acuity stimuli were rendered on the external display and imaged through the pupil relay to demonstrate that spatial frequencies up to 54 cycles per degree, corresponding to 20/11 visual acuity, are resolved. The completed external display was then used to render fixation markers across the field of view of the monitor, and a continuous retinal montage spanning 9.4 by 5.4 degrees of visual angle was acquired with the AOSLO. We conducted eye-tracking experiments during free-viewing and high-acuity tasks with polychromatic images presented on the external display. Sub-arcminute eye position uncertainty was achieved, enabling precise localization of the line of sight on the monitor while simultaneously imaging the fine structure of the human central fovea. This high refresh rate display overcomes the temporal, spectral, and field of view limitations of AOSLO-based stimulus presentation, enabling natural monocular viewing of stimuli in psychophysics experiments conducted with AOSLO.

## 1. Introduction

The adaptive optics scanning laser ophthalmoscope (AOSLO) has enabled high-resolution imaging of the human fovea [1–4]. Advances in the optical design and hardware for AOSLO systems have led to greater resolution and enhanced contrast for the smallest foveal cones, making it possible to investigate the anatomy of the living human retina *in vivo* [5–13]. In addition to high-resolution imaging, AOSLOs can be used to study fixational eye movements and conduct human psychophysics experiments. By projecting a stimulus directly onto the retina and collecting the backscattered light, it is possible to unambiguously locate the stimulus position on the retina. This procedure has been used to investigate fixation behavior and identify anatomical features of the human retina that may influence behavior [10,13–21]. Unlike other eye-tracking methods that rely on calibrations to accurately localize the line of sight and then transform that information into retinal image motion, AOSLO-based eye-tracking measures retinal motion directly [22–25].

Despite this spatial accuracy advantage of AOSLO systems, a number of constraints on visual stimulation limit their use in vision research. Stimuli are often rendered with monochromatic light at luminance levels that are much greater than natural viewing conditions. The field of view of an AOSLO is limited to the angular extent of the raster scan and is often selected to be around 1 degree by 1 degree for the full field of view to achieve sufficient pixel density for resolving the smallest foveal cones [10,13,18,20]. Such a small field of view limits the stimulus size to a maximum of 1 degree in width and precludes the presentation of stimuli outside the central fovea. Correction of the aberrations of the human eye using adaptive optics is necessary for resolving the smallest foveal cones, and this technique also enables scientists to study the impact of ocular aberrations by quantifying the improvement in visual acuity when aberrations are corrected [26–28]. However, this aberration correction results in a visual experience that is unnatural, where high spatial frequencies are amplified to contrast levels that are impossible to achieve in natural viewing because of blur from diffraction when the pupil is small (i.e., less than 3 mm diameter) and blur from aberrations when the pupil is large.

In addition, the temporal frequency of stimulus presentation is low. The scanning architecture of AOSLO means that the stimulus projected onto the retina is not temporally stationary but is composed of a focused spot of light that is rapidly scanned across the retina and modulated to produce a fixation marker or visual stimulus. The low frame rate of 30 Hz, which is standard for many AOSLOs, causes the stimulus and the raster pattern to flicker. Neurons in the retina and the early visual cortex can respond to higher temporal frequency ranges [29–31], and temporal modulations—which are not present when viewing a natural scene or looking at a high refresh rate LCD screen—are known to affect visual sensitivity in humans [32]. These temporal constraints contribute to the unnatural viewing conditions of AOSLO stimulation and may complicate the generalization of AOSLO psychophysics data to more natural conditions.

To overcome these hardware constraints, we present a custom high refresh rate display for use in AOSLO psychophysics experiments. An LCD monitor with a refresh rate of 360 Hz enables high temporal resolution and avoids the visual transients introduced by the 30 Hz AOSLO raster scan. The monitor supports full-color rendering of stimuli instead of monochromatic illumination. The full field of view is 45 times larger in area than the standard 1-degree AOSLO raster. This display is viewed through a relay that compensates for the spherical refractive error of the eye while leaving the other aberrations of the eye unchanged, and an external pupil with a diameter of 3 mm ensures that the stimulus is viewed through a natural pupil size instead of a fully dilated pupil. Eye clearance greater than 80 mm enables integration with a standard AOSLO.

Prior art has demonstrated external displays for use with AOSLO, but these displays have not achieved the requirements outlined above, which are necessary for natural viewing of stimuli. Putnam et al. utilized a digital micromirror device (DMD) to display a stationary fixation marker in an adaptive optics ophthalmoscope [33]. This DMD display was placed before the deformable mirror, meaning that the stimulus was viewed with ocular aberrations corrected. Aberration correction and monochromatic stimulus delivery at 550 nm created unnatural stimulus viewing conditions in this experiment, and the maximum frame rate of the DMD was not reported. Steven et al. implemented a full-color display for presenting a fixation marker, but at relatively low spatial resolution (i.e., 0.4 pixels per arcmin) and temporal resolution (i.e., 120 Hz maximum) [34]. Adaptive optics visual simulators and Tracking Scanning Laser Ophthalmoscopes (TSLOs) have utilized external displays that satisfy some of the requirements for natural viewing. However, these systems lack the ability to simultaneously provide rich visual stimulation while imaging the human retina with cellular resolution [26,35]. The external display reported in this paper achieves full-color stimulus presentation at high spatial and temporal resolutions without compensating for higher-order ocular aberrations, enabling natural monocular stimulus presentation in AOSLO psychophysics experiments. We demonstrate the capabilities of this external display integrated with an AOSLO by capturing high-resolution images of the retina during a fixation task, and by tracking eye movements during free-viewing and high-acuity tasks with stimuli rendered on the external display.

## 2. Optical design and specifications

### 2.1 Display selection and first-order layout of optical relay

We selected a high refresh rate monitor instead of a projector to satisfy the high spatial and temporal resolution requirements. There are only a few commercially available projectors with frame rates greater than 300 Hz, all of which have pixel sizes that are incompatible with the sampling resolution and pupil diameter requirements for projecting high-acuity visual stimuli (i.e., greater than 30% contrast at 30 cycles per degree) through an effective pupil diameter of 3 mm.

We placed the monitor approximately four meters from the eye pupil to achieve high pixel density (i.e., high spatial resolution). We then utilized an optical relay between the eye and the monitor to relay the finite object distance to real and virtual image distances corresponding to the far point for a given subject, making it possible for myopic and hyperopic individuals to see the monitor clearly. The second purpose of the relay is to create an effective pupil for the subject. During experiments with the AOSLO, the pupil of the eye is dilated to achieve sufficient numerical aperture for resolving the smallest foveal cones. Viewing high-acuity stimuli through a dilated pupil causes significant blur and reduced visual acuity because of ocular aberrations, especially spherical aberration. To overcome this challenge, an external iris is placed conjugate to the eye pupil and is adjusted to a diameter of 3 millimeters, which achieves a good balance between diffraction and ocular aberrations for the average human eye [36–39]. To satisfy these requirements, we designed a custom optical relay with unit magnification. Inspiration for the design was drawn from a variation of the Badal Optometer, which achieves nearly constant monitor pixel angle (i.e., spatial resolution of the monitor) over a large range of refractive error corrections for a distant monitor [40].

Other requirements for the custom optical relay are set by the space constraints. To enable integration with our AOSLO and sufficient clearance between the eye pupil and the dichroic mirror, the eye clearance was specified to be at least 80 mm. This specification can be achieved by choosing a lens system with a back focal length of at least 135 mm, which accounts for the space required to install a 2-inch diameter fold mirror between the eye and the relay. Fig. 1 shows a schematic of the external display and illustrates how the monitor, optical relay, and AOSLO are integrated. The field of view is determined by the size of the monitor and the distance between the monitor and the entrance pupil of the relay. The nominal full field of view at 0 diopters of correction is 9 x 5 degrees, which can be achieved with no vignetting using 2-inch diameter lenses and a 3-inch diameter fold mirror. The refractive error correction range was chosen to cover at least 10 diopters, with the mean value shifted toward myopic correction to account for the high prevalence of myopia worldwide and in the subject population [41]. A range of -8 diopters to +4 diopters was selected based on a balance between overall system length, travel range of the stage, and the eye clearance requirement.

**Fig. 1.**
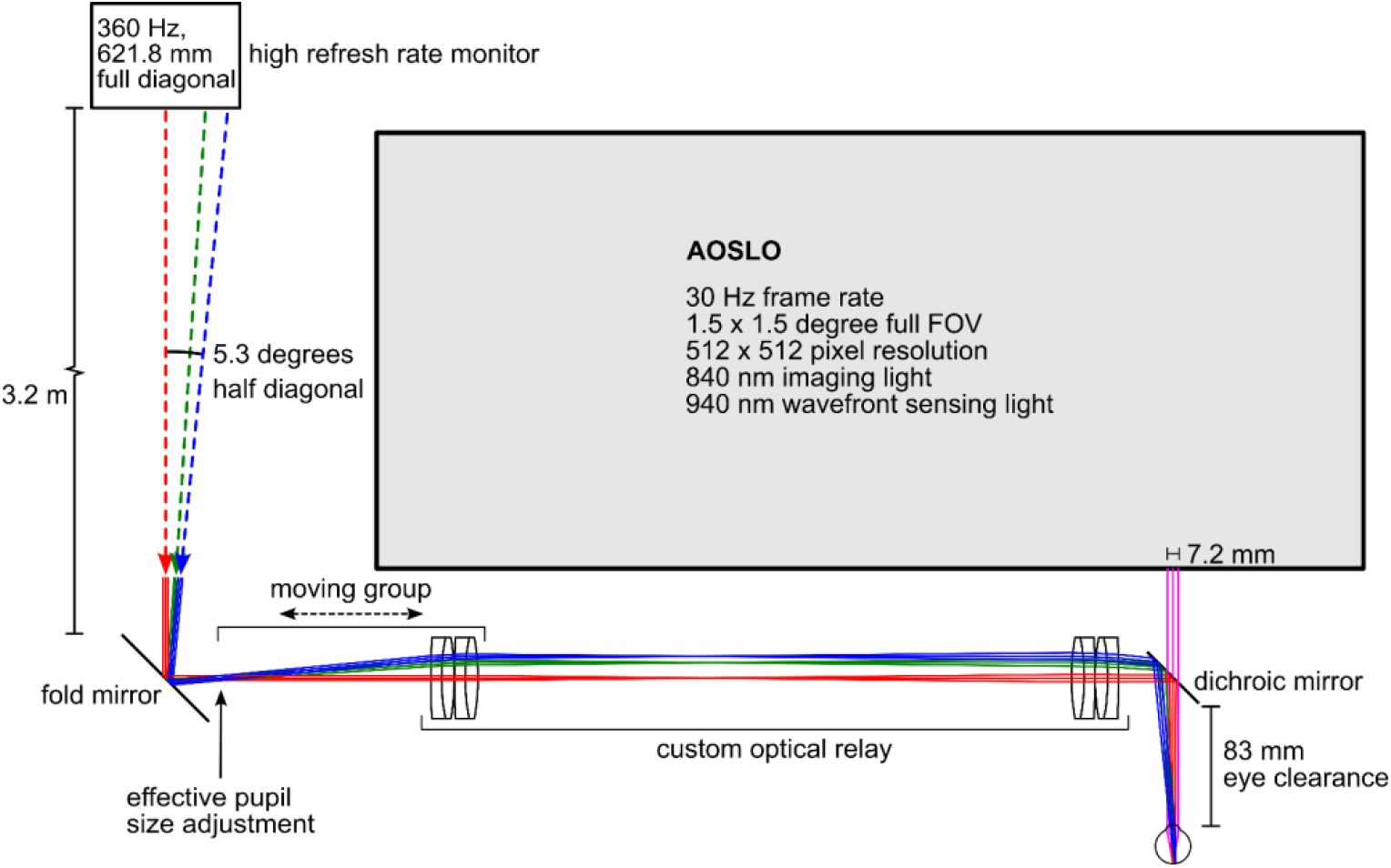
System layout showing the high-speed display integrated with our custom AOSLO [42]. A 7.2-mm diameter beam exits the AOSLO, consisting of 840 nm imaging light and 940 nm wavefront sensing light to measure and correct ocular aberrations. The AOSLO field of view is adjusted to its maximum value of 1.5 degrees to enable imaging and eye-tracking over a larger region of the retina compared with the standard 1-degree field of view. A 360 Hz monitor is positioned across the room from where the subject is seated. Light from the monitor is directed toward a fold mirror and then through the entrance pupil of a custom optical relay. After passing through the two lens groups of the optical relay, the visible light from the monitor is reflected off a dichroic mirror toward the eye. Imaging and eye-tracking, both working at 840 nm, are conducted with the AOSLO while visual stimuli are presented on the 360 Hz monitor and viewed through an external pupil with a diameter of 3 mm.

### 2.2 Specifications

Detailed specifications for the monitor and the relay are listed in Table 1. The monitor has sufficient pixel density to display high-acuity stimuli up to the Nyquist limit of 107 cycles per degree, which is much higher than the sampling limit imposed by the cone mosaic of approximately 60 cycles per degree [10,17,27,43,44]. The two lens groups that make up the optical relay have effective focal lengths of 153 mm and back focal lengths of 139 mm. This allows for the installation of a 2-inch flip mount fitted with a dichroic mirror between the eye and the fixed lens group. The eye clearance—measured from the edge of the flip mount to the eye pupil—is 83 mm. The performance target for the design is better than 0.07 waves RMS wavefront error across the full field of view and the whole correction range from -8 diopters to +4 diopters.

**Table 1:** Specifications for the monitor and optical relay that make up the high-speed display.

| Description | Target | Actual | Notes |
| --- | --- | --- | --- |
| *Monitor specifications* |  |  |  |
| Monitor width (mm) | 543.17 | Same | Full width of active area |
| Monitor height (mm) | 302.62 | Same | Full height of active area |
| Resolution (pixels) | 1920 x 1080 | Same |  |
| Pixel dimensions (mm) | 0.2829 x 0.2802 | Same | Pixel width and height respectively |
| Refresh rate (Hz) | $\geq 300$ | 360 | |
| *Relay specifications* |  |  |  |
| Horizontal full field of view (degrees) | $\geq 8$ | 9.0 | Measured at eye pupil at 0-diopter configuration |
| Vertical full field of view (degrees) | $\geq 4$ | 5.0 | Measured at eye pupil at 0-diopter configuration |
| Effective pupil diameter (mm) | 3.0 | Same | Chosen for balance between diffraction and aberrations in the human eye |
| Wavelengths (nm) | 486.1, 587.6, 656.3 | Same | Visible spectrum with equal weighting in optical design software |
| Refractive error correction range (diopters) | -8 to +4 | -8.02 to +4.00 | Limited by travel range of the motorized stage |
| Monitor angular resolution (pixels/arcmin) | $\geq 3$ | 3.6 | At 0-diopter configuration. Corresponds to the Nyquist frequency of 107 cycles/degree |
| *Performance specifications* |  |  |  |
| Maximum RMS wavefront error (waves at 543 nm) | $\leq 0.07$ (design)<br>$\leq 0.114$ (as-built) | 0.048 (design)<br>0.106 (as-built) | Maximum error across all configurations and field points |
| Contrast at 30 cycles per degree (%) | $\geq 30$ | 44.7 | Contrast was measured at 36 cycles per degree due to pixel discretization |
| Distortion (%) | $\leq 1.0$ | 0.08 | Modeled in optical design software |
| *Mechanical constraints* |  |  |  |
| Monitor distance (m) | $\geq 4$ | 4.1 | Measured from eye pupil to monitor |
| Eye clearance (mm) | $\geq 80$ | 83.1 | Distance from edge of flip mount to eye pupil |
| Travel range of moving group (mm) | $\leq 300$ | 279.4 | Total translation of moving group from negative extreme to positive extreme |
| Maximum element diameter (mm) | $\leq 76.2$ | Same | All doublets are 2-inch in diameter. Fold mirror is 3-inch in diameter |

The as-built performance target is driven by the need to display high-acuity visual stimuli and is set to be greater than 30% contrast at 30 cycles per degree (i.e., the fundamental spatial frequency of the 20/20 line of the eye chart). This criterion was validated through modeling in CODE V optical design software (Sunnyvale, California), where we determined, by examining the polychromatic Modulation Transfer Function (MTF), that the optical relay maintains greater than 30% contrast at 30 cycles per degree for Strehl ratios greater than 0.6. Since we validated our assembly using a Shack-Hartmann wavefront sensor, it is useful to express the MTF performance and Strehl ratio in terms of the RMS wavefront error. Using equation 3 in [45], we find that a Strehl Ratio of 0.6 corresponds to 0.114 waves RMS wavefront error. Therefore, the as-built performance is set to be no worse than 0.114 waves (i.e., ≤ 0.114) RMS wavefront error across the full field of view and correction range, as shown in Table 1.

The maximum RMS wavefront error of 0.048 waves for the design was determined through modeling in optical design software, and the maximum as-built RMS wavefront error of 0.106 waves was measured with a wavefront sensor, which is described in Section 4.1.

### 2.3 Optical design and performance assessment

After determining the specifications for the optical relay, we designed the optical system using off-the-shelf lenses and mirrors. Achromatic doublets were selected to provide color correction over the range of visible wavelengths. We selected doublets with the same focal lengths to achieve unit angular magnification. Since data from our optical design simulations indicated that two identical doublets were not sufficient to meet the performance targets, we proceeded with a design that utilized four identical doublets (ACT508-300-A from Thorlabs), with two lenses in each group. In the optical design software, the total distance between the monitor and the eye was fixed. A lens module with a fixed focal length of 16.67 mm and an image plane distance that varied through zoom was used to simulate an eye with varying amounts of refractive error. The axial position of the moving lens group was allowed to vary through zoom to compensate for the refractive error of the model eye. Optimization was conducted with all lens variables frozen and only the relevant airspaces varying, while also constraining the total distance between the monitor and the eye. Fig. 2 shows the lens drawing across the correction range, which demonstrates how the moving group shifts to compensate for refractive errors.

**Fig. 2.**
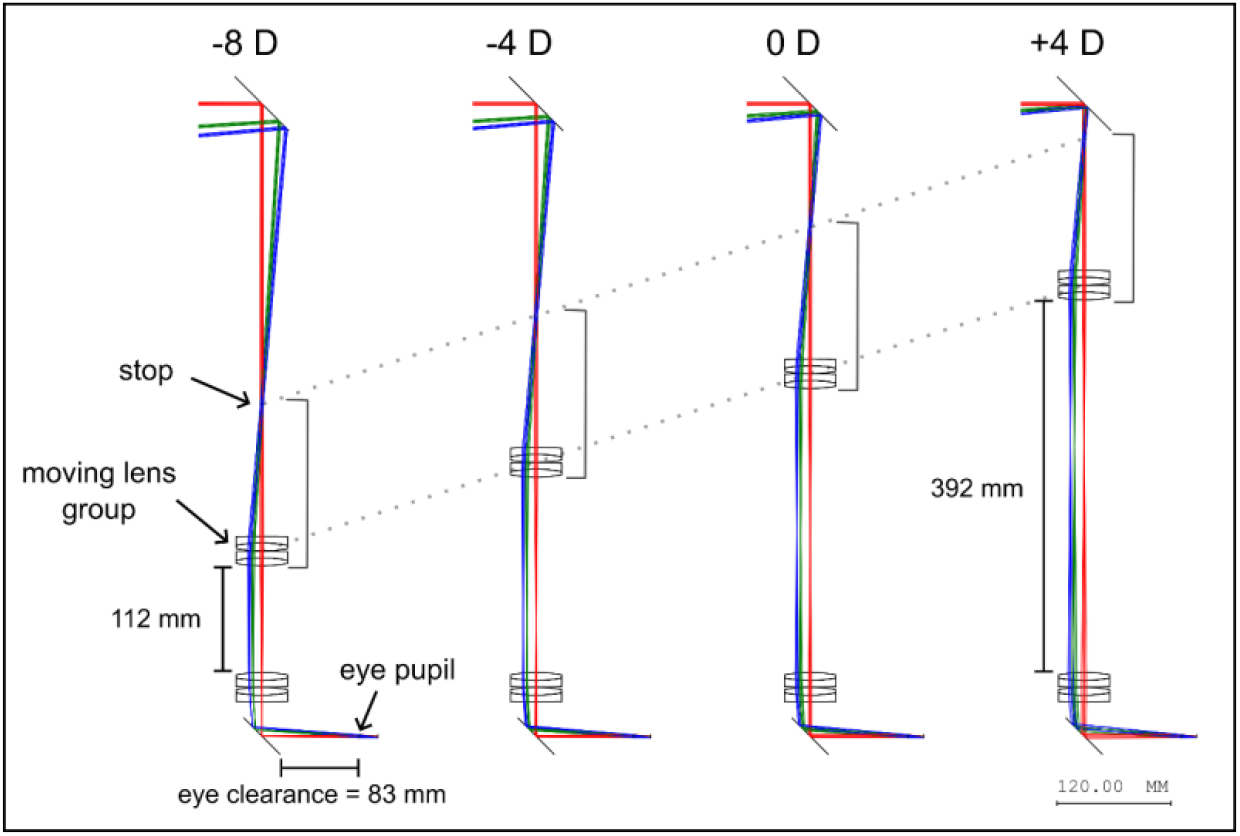
Lens drawing of the optical relay showing the motion of the moving group across the full correction range from -8 diopters to +4 diopters. The lenses and the aperture that make up the moving group are attached to a lens tube, and the whole assembly moves together on a motorized translation stage. The total travel range of the moving group is 280 mm (11 inches). The eye clearance from the edge of the dichroic mirror to the eye pupil is 83 mm.

Performance of the optical design was assessed using RMS wavefront error and MTF. The maximum RMS wavefront error of 0.0482 waves occurred at the corner of the field of view for the +4-diopter configuration. The design meets the diffraction-limited criterion of RMS wavefront error less than 0.07 waves across the full field of view and correction range. The MTF was evaluated to determine the contrast at 30 cycles per degree, which corresponds to the fundamental spatial frequency of the 20/20 line of an eye chart. To convert between length and angular units at the image plane, the retinal magnification factor for a 60-diopter model eye was used, which results in a conversion factor of 0.291 mm per degree of visual angle. The dashed horizontal and vertical lines in Fig. 3 show the MTF at 30 cycles per degree, or 103 cycles per mm. The MTF is greater than 0.45 at 30 cycles per degree across the full field of view and the full correction range. The maximum distortion is less than 0.1%. These results suggest that the design can render high-acuity stimuli across the full extent of the monitor with high contrast and low distortion.

**Fig. 3.**
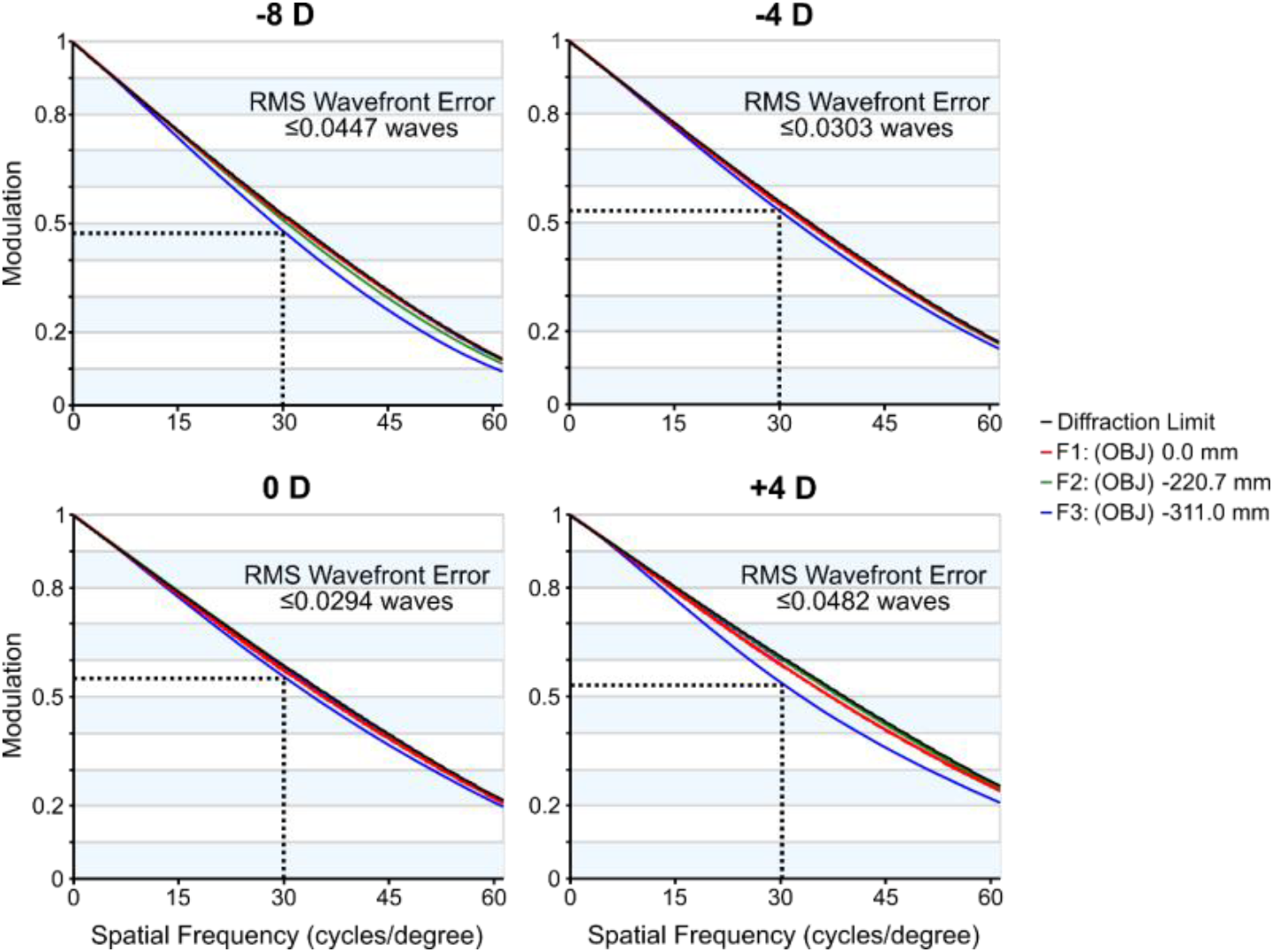
Performance evaluation for the optical design of the custom optical relay. Polychromatic MTF curves are shown for different refractive error correction settings. The dashed horizontal and vertical lines show the MTF value at 30 cycles per degree, which is greater than 0.45 for all fields and configurations. The maximum RMS wavefront error for each configuration is also reported, and all values are better than the performance target of 0.07 waves for the design.

## 3. Optomechanical design and assembly

### 3.1 Constructing a CAD model for the optical relay

The finished optical design was exported as a CAD model from the optical design software and imported to Onshape (Boston, Massachusetts) for the optomechanical design. Lens mounts and holders were added to the model from the Thorlabs catalog. A motorized stage from Velmex, Inc. (Bloomfield, NY) was selected for its compact size, 12-inch travel range, vertical mounting compatibility, and high straight-line accuracy. The stage was incorporated into the optomechanical model and custom adapter plates were designed to interface between the stage and other components. A construction rail with a length of 1 meter served as the reference plane and attachment point for all components. The completed CAD model is shown in Fig. 4A.

**Fig. 4.**
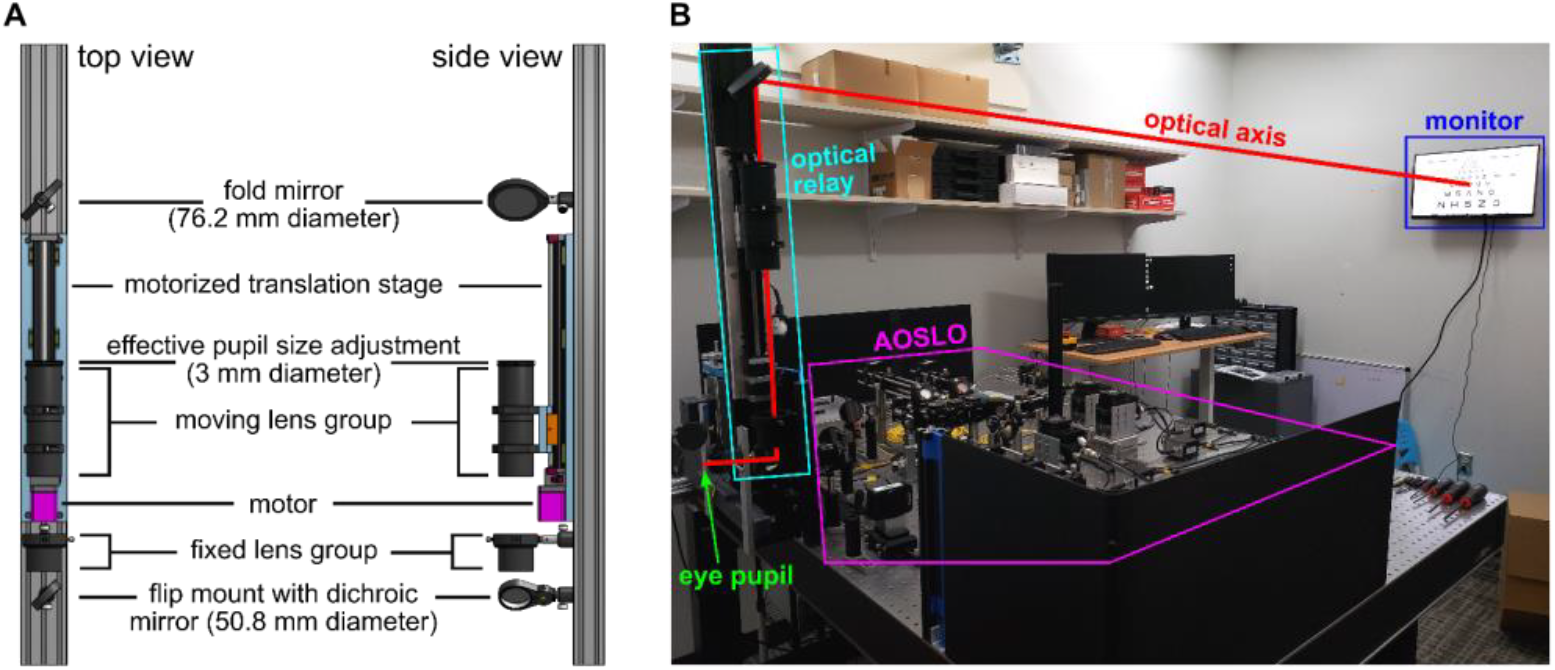
Mechanical layout of the external display components. A. A CAD model shows the custom optical relay design. A moving lens group, which includes a 3-mm diameter aperture stop, translates vertically on a motorized stage. A dichroic mirror combines the high-speed display with the AOSLO and is mounted on a precision flip mount, enabling the high-speed display to be quickly added or removed from the full system. B. The external display after integration with the AOSLO. The optical axis for the custom optical relay is traced from the monitor to the eye pupil to demonstrate the light path.

### 3.2 Assembly and alignment

Using the CAD model, we tabulated all the component positions relative to the end of the construction rail and used these measurements for assembly of the optical relay. We placed an alignment laser (PL201 from Thorlabs) at the far end of the construction rail, beyond the post location for the fold mirror mount shown in Fig. 4A. The fold mirror and dichroic mirror were not installed during the initial alignment to facilitate ease of alignment and testing of the optical relay. Near and far alignment guides (SM1A7 from Thorlabs) were used to center the alignment laser beam at both ends of the construction rail and ensure that the beam was parallel to the reference plane defined by the front surface of the construction rail. An empty lens tube was then attached to the motorized stage to facilitate coalignment of the stage with the construction rail. Ground glass alignment disks with a central hole diameter of 2 mm (DG20-1500-H2-MD from Thorlabs) were used to locate the center of the lens tube. The stage was aligned to keep the laser beam centered on the lens tube as the stage translated between the two ends of its range of motion, enabling coalignment within +/-0.5 mm tolerances when used with the alignment laser with a beam diameter of 3 mm. The fixed lens group utilized a mechanical mount with precision x-y translation, which enabled precise alignment of the lens tube used in this group, achieving the same +/-0.5 mm alignment tolerances as the other lens tube.

After the mechanical components were installed and aligned, the lenses were installed into their respective lens tubes. A single retaining ring with a width of 2.5 mm was used to separate the two lenses in each group. This configuration achieves a vertex-to-vertex airspace of 0.18 mm between the two lenses, which corresponds to the airspace used in the optical design software. After the lenses were installed in the lens tubes, the alignment was checked using the alignment laser and the ground glass alignment disks, which were placed at the front and back of the moving lens group and the back of the fixed lens group. Alignment was maintained within the +/-0.5 mm beam centration tolerances after the lenses were installed.

## 4. Testing and calibration

### 4.1 Wavefront measurements across the field of view and correction range

To test the performance of the assembled optical relay and to calibrate the refractive error correction settings, we placed a Shack-Hartmann wavefront sensor (WFS40-7AR from Thorlabs) at the exit pupil. A well corrected input wavefront was prepared by collimating the output of a single-mode fiber using an achromatic doublet with a focal length of 40 mm. An iris with a diameter of 3 mm was placed one focal length beyond the collimating lens to set the entrance pupil diameter. Shack-Hartmann measurements of the input wavefront showed an RMS wavefront error of 0.033 waves, which is well below the diffraction limit (i.e., less than 0.07 waves). The wavefront tests were conducted with 543 nm light, which is near the center of the visible spectrum. A single wavelength was used for wavefront testing because the design is well corrected for chromatic aberrations. Astigmatism, field curvature, and coma are the limiting aberrations across the whole correction range based on modeling in optical design software.

To calibrate the refractive error correction across the range of motion of the motorized stage, the Shack-Hartmann wavefront sensor was used to measure the defocus introduced by the optical relay at 15 different refractive error settings, corresponding to a sampling interval of 0.8 diopters. The optical relay has a linear relationship between the separation of the lens principal planes and the introduced defocus, similar to the Badal Optometer [40], so the points were fit to a line to generate the calibration between the motorized stage position and the defocus. This calibration was then implemented in a custom MATLAB (Natick, Massachusetts) application for controlling the motorized stage and setting the refractive error correction. The full correction range was found to be -8.02 diopters to +4.00 diopters, which agrees well with the results from optical modeling.

Measurements of the residual wavefront error were first collected for the on-axis field point at four defocus settings across the refractive error correction range using the Shack-Hartmann wavefront sensor. The wavefront maps were represented in a Zernike polynomial basis with the first 16 terms in the FRINGE definition using ZernikeCalc [46]. The first three FRINGE Zernike terms—piston, x-tilt, and y-tilt—were subtracted from the wavefront maps because these terms do not impact image quality. For the corrections at -8 diopters, -4 diopters, and +4 diopters, the defocus term (i.e., Z4) was also subtracted from the wavefront. Subtracting the defocus from these three measurements was necessary because a large amount of defocus was purposely introduced by the optical relay at these positions to correct for ocular refractive errors, and this defocus must be removed before quantifying the residual wavefront aberrations. After subtracting these Zernike polynomials from the wavefront maps, the residual wavefront errors were quantified and reported in Table 2. The RMS wavefront error ranged from 0.032 waves at +4 diopters to 0.059 waves at -8 diopters for the test wavelength of 543 nm.

**Table 2:** RMS and peak-to-valley wavefront error for the on-axis wavefront measurements across the correction range.

| Refractive error correction (diopters) | -8 | -4 | 0 | +4 |
| --- | --- | --- | --- | --- |
| RMS wavefront error (waves at 543 nm) | 0.059 | 0.043 | 0.033 | 0.032 |
| Peak-to-valley wavefront error (waves at 543 nm) | 0.296 | 0.212 | 0.203 | 0.206 |

After validating the on-axis performance, wavefront measurements were collected across the full field of view of the optical relay. The entrance pupil of the collimated input was displaced laterally and then tilted to center the beam on the entrance pupil of the optical relay for each measurement point. The wavefront sensor remained positioned at the exit pupil of the relay for all measurements. A set of three repeated measurements was collected at each of 17 field points spanning the full field of view. Fig. 5 shows how the 17 measurement points were distributed across the field of view. To analyze the wavefront performance, each set of three measurements was first averaged and the standard deviation was computed to ensure repeatability of the measured wavefront. The average standard deviation of the wavefront over the full pupil was 0.0033 waves (range of 0.0018 to 0.0052) for the 17 different measurements, which demonstrates good repeatability across each set of repeated measurements.

**Fig. 5.**
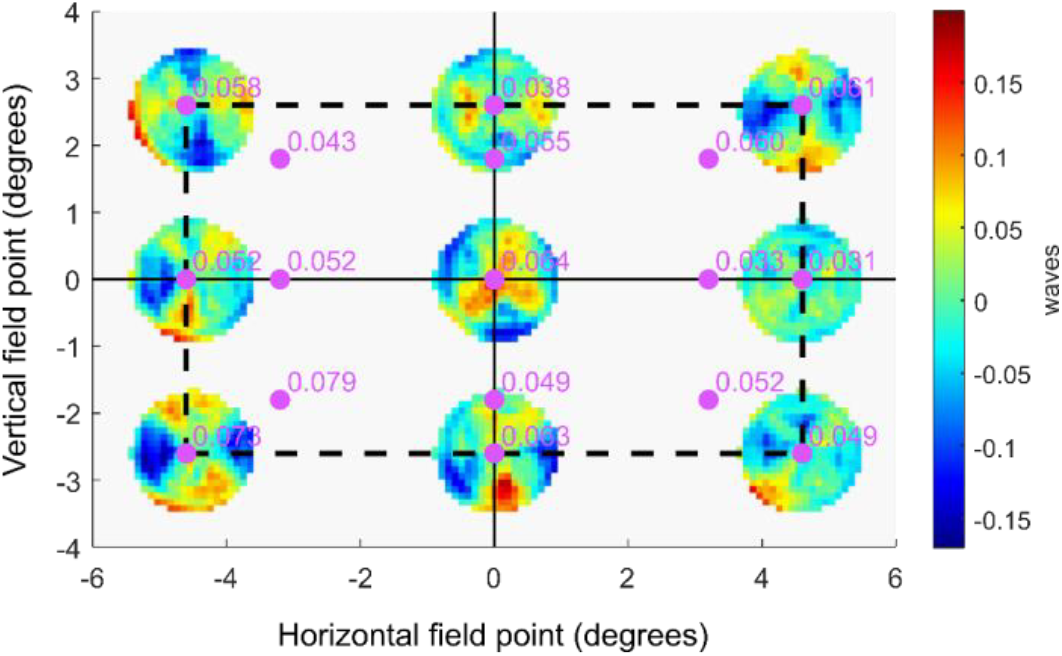
Wavefront measurements across the field of view for the 0-diopter configuration. Measurements were collected at 17 different points across the field of view. The reported values are the RMS wavefront error after subtracting the field-averaged residual focus and the piston, tip, and tilt for each field point measurement. The RMS wavefront error satisfies the diffraction limit for the horizontal and vertical meridians across the full field of view, but some of the diagonal points are slightly above the diffraction-limited criterion, while still meeting the as-built performance target of less than 0.114 waves. Over the full field of view, the maximum RMS wavefront error is 0.079 waves, which occurs toward the lower left corner of the field of view. The average RMS wavefront error is 0.054 waves. The units are waves at 543 nm, which is the wavelength used for testing.

The average wavefront maps were then examined as described above. Because the optical design has a small amount of residual field curvature, the average defocus term (i.e., Z4 in FRINGE definition) was computed across the 17 field point measurements and was then subtracted from each of the average wavefront maps. Subtracting the field-averaged defocus is beneficial for assessing the performance across the field of view, and it enables direct comparison with wavefront analysis results from optical design software, which automatically compensate for the average defocus across the field of view. Following the same procedure used for the on-axis wavefront measurements, the Zernike terms for piston, x-tilt, and y-tilt were then subtracted from the average wavefront maps before quantifying the residual RMS wavefront error.

The resulting wavefront error maps for 9 of the 17 field points are shown in Fig. 5 for the 0-diopter configuration. RMS wavefront errors are reported on the figure for each of the 17 measurement points. The average RMS wavefront error across the full field of view is 0.054 waves, which is below the diffraction limit. The maximum wavefront error of 0.079 waves occurs in the lower left portion of the field of view. This maximum value is within the as-built tolerance of less than 0.114 waves as reported in Section 2.2. The discrepancy between the on-axis measurements reported in Table 2 and Fig. 5 (i.e., 0.033 waves and 0.064 waves respectively) comes from the different methods used to compensate for defocus in the two measurements. For the on-axis measurement reported in Table 2, defocus was optimized for the on-axis field point. The results shown in Fig. 5 use the field-averaged defocus to represent average performance across the field of view, which introduces some defocus to the on-axis field point to achieve better average correction across the field of view.

Wavefront measurements across the field of view were repeated at three other refractive error correction configurations, spanning the whole range from -8 diopters to +4 diopters, following the same procedures described above. Results are summarized in Table 3. The performance was best at +4 diopters, where the field-averaged RMS wavefront error was 0.048 waves, and the maximum was 0.07 waves. The worst performance occurred at -8 diopters, which had a field-averaged RMS wavefront error of 0.077 waves and a maximum RMS wavefront error of 0.106 waves. These results demonstrate that the as-built performance is within the RMS wavefront error specification of less than 0.114 waves across the whole field of view and correction range. The corresponding as-built Strehl ratio target of greater than 0.6 is also satisfied.

**Table 3:** Performance metrics across the field of view and correction range.

| Refractive error correction (diopters) | -8 | -4 | 0 | +4 |
| --- | --- | --- | --- | --- |
| Average RMS wavefront error (waves at 543 nm) | 0.077 | 0.060 | 0.054 | 0.048 |
| Maximum RMS wavefront error (waves at 543 nm) | 0.106 | 0.079 | 0.079 | 0.070 |
| Minimum Strehl ratio across the field of view | 0.64 | 0.78 | 0.78 | 0.82 |

### 4.2 Coalignment of the display and the AOSLO

After completing the wavefront measurements, the dichroic mirror was installed, and the optical relay was integrated into the AOSLO. The alignment laser that was used during assembly and alignment of the lenses and the optomechanics—described in Section 3.2—was reinstalled and aligned to pass through the center of each lens. This alignment laser was then used to identify and co-align the optical axis of the relay with the optical axis of the AOSLO. Near and far alignment guides—separated by 705 mm along the optical axis—were centered on the 680 nm AOSLO beam which was transmitted through the dichroic. The green alignment laser was then turned on and small adjustments to the position and angle of the optical relay were carried out until the alignment laser—which reflected off the dichroic—was centered on both the near and far alignment guides within the +/-0.5 mm alignment tolerances.

We used a monochrome camera with 2592 x 2048 pixel resolution and 4.8 μm pixel pitch (part number 34-851 from Edmund Optics) to conduct pupil matching between the AOSLO and the optical relay. The image sensor was placed at the eye pupil plane of the AOSLO, which was found by moving the camera axially until the edge of the deformable mirror aperture was in focus. This also corresponds to the position where the AOSLO laser beam appears stationary in time because the scanners are conjugate to the eye pupil plane. Next, the AOSLO laser was turned off and a flashlight was used to illuminate the entrance pupil of the optical relay. The axial position of the camera was adjusted until the edge of the iris at the entrance pupil was in best focus. The axial shift of the camera was then used to determine the distance by which the optical relay needed to move to achieve axial pupil matching. After making this small adjustment to the axial position of the optical relay relative to the AOSLO, measurements were conducted to verify that the two subsystems remained co-aligned.

At the eye pupil plane, the camera image sensor was used to quantify the overlap between the AOSLO beam and the alignment laser. The centers of the two beams were aligned within +/-0.3 mm in both the x- and y-directions. In the far plane, a ruler was used to verify that the beam centers were aligned within +/-0.5 mm in both the x- and y-directions. These alignment tolerances correspond to a maximum angular misalignment of 5.5 arcmin between the two subsystems. This maximum angular misalignment corresponds to an offset of 20 pixels on the monitor, which is approximately 1% of the monitor width. For axial pupil matching, the two subsystems are aligned within +/-5.2 mm, based on the depth of focus for the last relay telescope in the AOSLO. These measurements confirm that the two systems are well aligned to each other.

Finally, the alignment laser was placed at the eye pupil plane and directed through the optical relay toward the wall where the high refresh rate monitor would be mounted. The 3- inch fold mirror was installed and aligned to make the beam parallel to the floor—and perpendicular to the wall—in the room. The beam position was marked on the wall and was then used to install the monitor at the correct position, with the center of the monitor being aligned to the optical axis of the optical relay. The assembled and integrated external display is shown in Fig. 4B.

### 4.3 Assessing image quality

The image quality of the completed system was validated by placing the monochrome camera described above, fitted with a fixed-focal-length lens, at the eye pupil plane and imaging a custom test stimulus displayed on the monitor through the optical relay. The lens has an effective focal length of 35 mm, a variable aperture from F/1.4 to F/16, and distortion less than 0.05% (part number 63-247 from Edmund Optics). The test stimulus was designed to test spatial frequencies between 14 cycles per degree and 107 cycles per degree, which is the Nyquist limit of the monitor. The test stimulus was composed of seven lines, corresponding to the seven spatial frequencies tested, with each line containing six Snellen E optotypes in various orientations. We placed the test stimulus in each of the four corners and the center of the monitor to enable image quality assessment over the full field of view. Pupil matching of the camera and the optical relay was achieved by stopping the camera lens down to its minimum aperture and then translating the camera in three dimensions until the camera image achieved maximum brightness. The camera was focused at infinity, and the optical relay was set to the 0-diopter configuration for this test.

Fig. 6A shows the test stimulus image over the full field of view of the monitor. A magnified view showing the lines with the smallest features is presented in panel B. Line 5 is resolved, while line 6 is not resolved: three distinct bars are visible for line 5, while the three bars are not visible for line 6. Line 5 was also resolved in the four corners of the monitor. Pixel gray value lineouts are shown for each of the six optotypes on line 5 in subpanels C – E. Line 5 has a stroke width of 2 pixels on the monitor, corresponding to 0.56 arcmin, and a fundamental spatial frequency of 54 cycles per degree. This spatial frequency approaches the human photoreceptor sampling limit of 60 cycles per degree. The average Michelson contrast for line 5 was 15% for both the horizontal and vertical bars. Results from this test demonstrate that the external display can present high-acuity stimuli up to the sampling limits imposed by the photoreceptor mosaic.

**Fig. 6.**
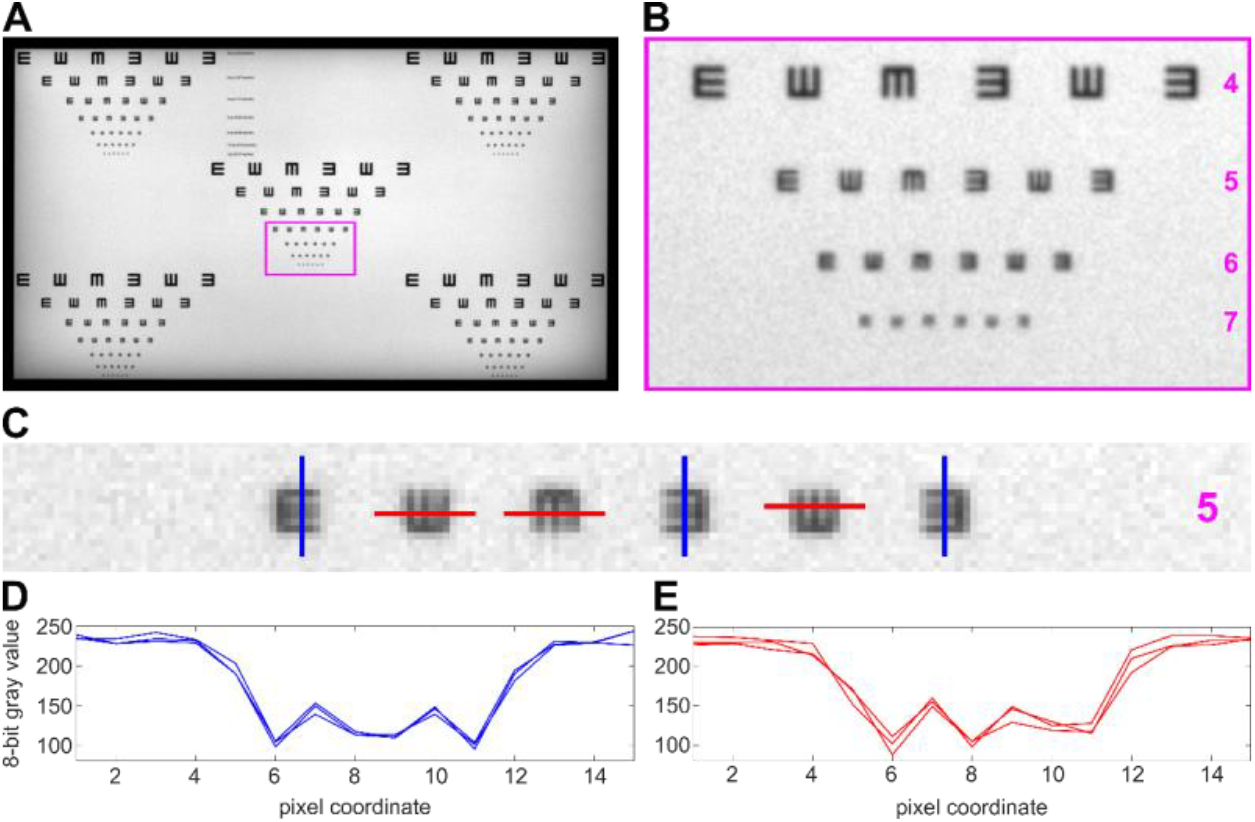
Resolution measurement of the assembled and aligned optical relay. A. A custom tumbling E test stimulus was designed and displayed on the monitor. It was then imaged with a fixed-focal-length camera. The test stimulus was placed in the center and each of the four corners of the monitor to assess resolution over the full field of view. B. Magnified view of the smallest features in the test stimulus, showing that row 5 is resolved. C. Further magnified view of row 5 with lines drawn to show where the pixel value lineouts were sampled. D. Pixel value lineouts for the three vertical lines. E. Pixel value lineouts for the three horizontal lines.

In addition to measuring the contrast of line 5, which is the last line to be resolved, the contrasts of lines 3 and 4 were measured across the whole image to quantify the resolution over the full field of view, as reported in Table 4. Line 3 has a stoke width of 4 pixels on the monitor, or 1.11 arcmin, with a spatial frequency of 27 cycles per degree. The features on this line are closest in size to the 20/20 line of the eye chart (i.e., 1.0 arcmin stroke width and 30 cycles per degree), subject to the pixel discretization of the monitor. There was not a significant difference between horizontal and vertical contrast measurements, and the mean values differed by less than 1%. The small standard deviations (i.e., less than 4.5% contrast) demonstrate that the image quality is consistent across the full field of view of the monitor.

**Table 4:** Measured contrast from the image quality assessment.

| Line | Stroke width (arcmin) | Spatial frequency (cycles/degree) | Snellen Acuity | Measured contrast (%) |
| --- | --- | --- | --- | --- |
| 3 | 1.11 | 27 | 20/22 | 64.2 +/- 3.2 |
| 4 | 0.83 | 36 | 20/17 | 44.7 +/- 3.0 |
| 5 | 0.56 | 54 | 20/11 | 15.4 +/- 4.2 |

## 5. Validation of human retinal imaging and eye-tracking

### 5.1 Retinal imaging over the full field of view of the display

The display was used to render fixation markers across the field of view of the optical relay, enabling human retinal imaging over an area spanning 9.4 by 5.4 degrees of visual angle. One individual with low myopia (−1 diopter), no known visual pathologies, and an age of 28 years participated in the human retinal imaging experiment. All procedures adhered to the ethical standards of the Research Subjects Review Board (RSRB) at the University of Rochester, which approved this study. The subject’s right eye was dilated with one drop each of 1% tropicamide and 2.5% phenylephrine ophthalmic solutions 15 minutes before the imaging session. The subject’s head was immobilized using a dental impression bite bar, and the subject’s pupil was aligned to the AOSLO exit pupil using a 3-axis translation stage.

The AOSLO used to capture the retinal images has been described in prior art [42]. The imaging full field of view was set to 1.5 by 1.5 degrees. A wavelength of 840 nm was used for imaging, and wavefront sensing was accomplished using 940 nm light and a custom Shack-Hartmann wavefront sensor. A deformable mirror with 97 actuators enabled wavefront correction over a 7.2 mm pupil diameter. The power levels for the two wavelength channels were 144 μW for 840 nm and 216 μW for 940 nm, which are both less than 12% of the maximum permissible exposure defined by the ANSI Z136.1 standard for an exposure duration of two hours [47].

A white square with an angular subtense of 5 arcmin was displayed on a black background on the monitor and served as a fixation marker. This fixation marker was placed at each of 45 different positions on the monitor during the experiment, spanning 8 degrees horizontally and 4 degrees vertically with a spacing of 1 degree. Given that the fixation markers were spaced by 1 degree from each adjacent fixation marker, images acquired at these fixation points were overlapping by approximately 33% in area for the 1.5-degree square imaging field of view. At least two videos were recorded with the AOSLO at each fixation point, and each recording had a duration of 10 seconds. The video frame rate is 30 frames per second.

The videos from each fixation point were then manually graded to identify the best video recorded at each fixation point, based on the sharpness of the image and fixational stability during the recording. The best 45 videos—one from each fixation point—were then registered using strip-based video registration to yield high-resolution and high-contrast image patches of the photoreceptor mosaic [48]. These image patches were then automatically montaged using open-source software [49] to generate a continuous image of the photoreceptor mosaic spanning 9.4 by 5.4 degrees of visual angle, or 2.8 by 1.6 mm on the retina using Bennett’s method to convert between angular and linear units based on the subject’s axial length [50]. Finally, the aligned image patches were loaded into CorelDRAW Photo-PAINT (Ottawa, Ontario) for manual layering and stitching. During this manual procedure, image patches with the highest sharpness for a given retinal region were brought to the foreground and the brightness was adjusted for several image patches to achieve uniform brightness across the montage. Stitch line visibility was reduced by feathering some of the sharp edges of the image patches using the eraser tool in Photo-PAINT. After merging the layers, the image was contrast enhanced using adaptive histogram equalization [51] to produce the result shown in Fig. 7.

**Fig. 7.**
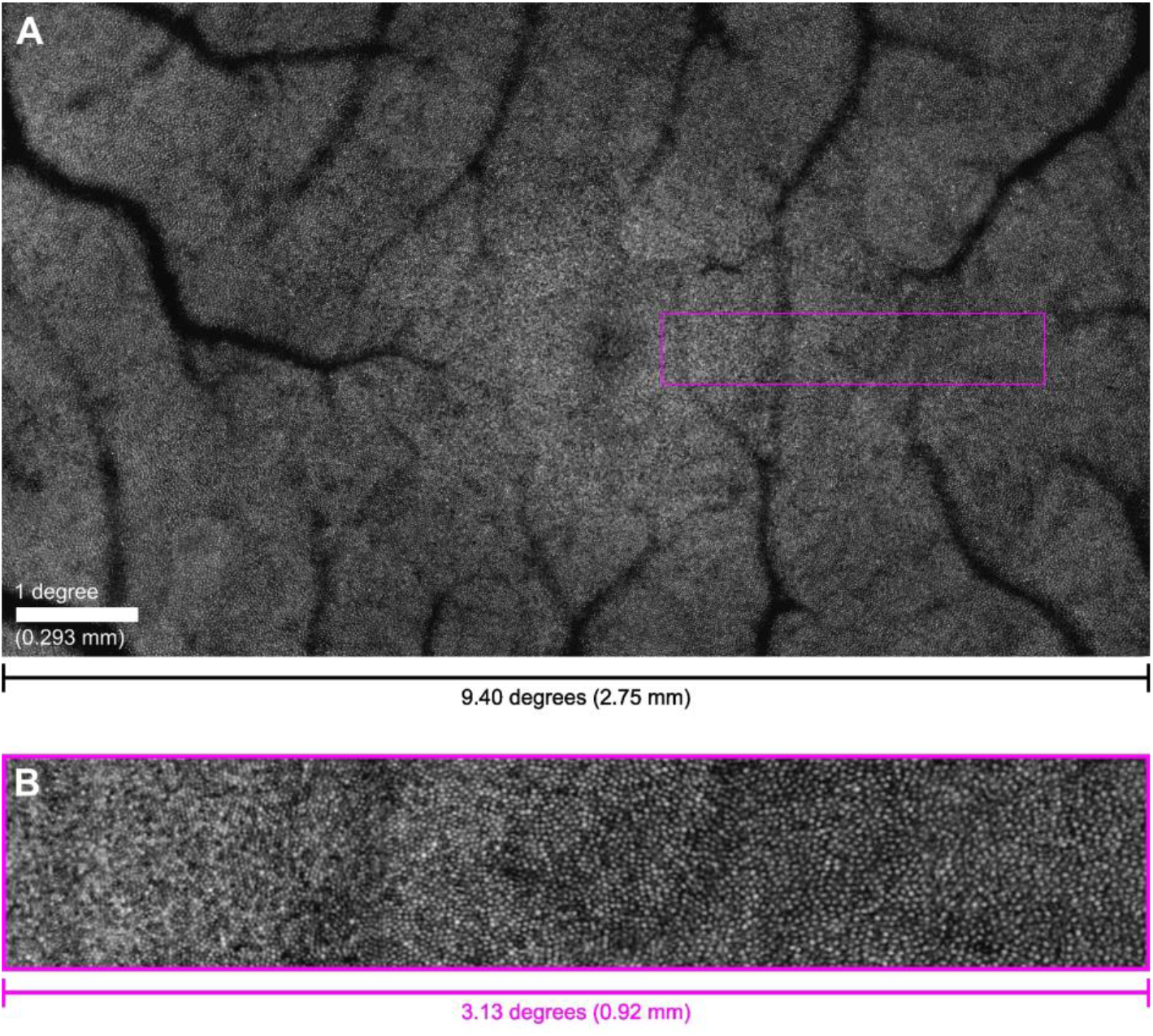
Full-field composite retinal image of a healthy human retina captured with the AOSLO using 840 nm imaging light and visible fixation markers presented across the full field of view of the display. A. The image spans 9.4 x 5.4 degrees of visual angle and encompasses the entirety of the 5-degree fovea. The dark region at the center of the image is the foveola, where cones are smallest and most densely packed. Superior retina is oriented up and nasal retina is to the right in the image. The subject’s right eye was imaged. B. Magnified inset (3x magnification) of the nasal retina showing the variation in cone size and packing from the edge of the foveola at the left side of the inset (0.43 degrees eccentricity) into the parafovea at the right side of the inset (3.56 degrees of eccentricity).

### 5.2 Offline eye-tracking using strip-based video registration

The integrated display and AOSLO system was also used to conduct retinal image-based eye-tracking in both a free-viewing task and a high-acuity task. The same participant from Section 5.1 completed these eye-tracking experiments. A subjective refraction was performed by iteratively adjusting the motorized optometer by 0.25-diopter steps in both directions from the expected refractive error setting while the participant viewed an eye chart. The subjective refraction resulted in a correction of -1 diopter, which matched the individual’s refractive error of -1 diopter as measured by an autorefractor.

Before collecting experimental data, we implemented an alignment procedure to register the coordinates of the AOSLO with the monitor coordinates. This procedure was necessary for mapping the eye-tracking results onto the images that were displayed on the monitor during the tasks. First, a 5-arcmin black square was presented at the center of the AOSLO raster scan by modulating the 840-nm imaging light with an acousto-optic modulator. The AOSLO raster was easily distinguished from the background when the monitor pixel values were set to their minimum value, and the edges of the black square were well defined because of the adaptive optics correction. The subject then used a keypad to move the position of a 3-arcmin gray square that was rendered on the monitor until the gray square was precisely centered inside of the black square, as shown in Fig. 8A.

**Fig. 8.**
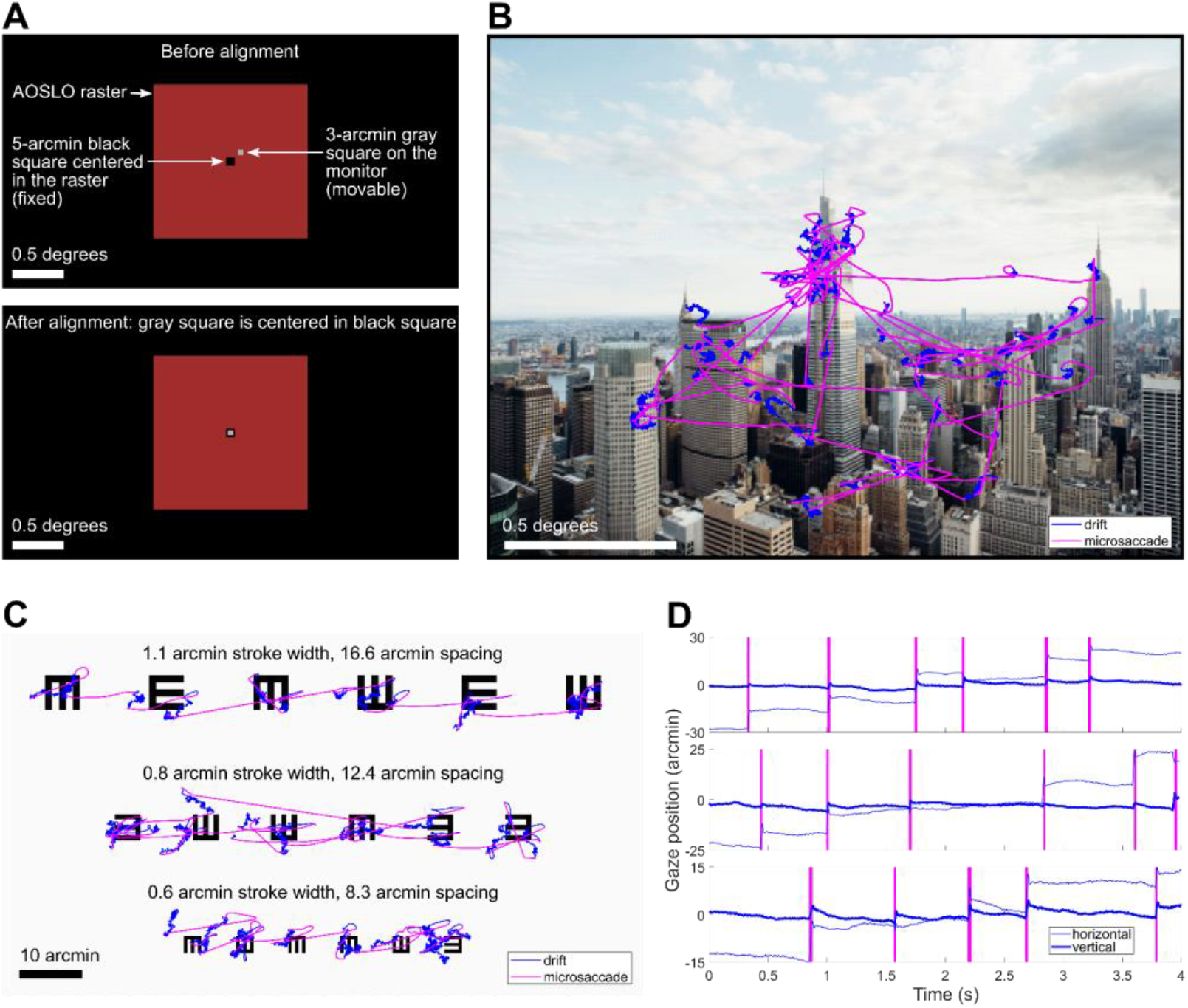
Results from the eye-tracking tasks conducted with the combined AOSLO and external display. A. Illustration of the alignment procedure for registering the monitor coordinates to the center of the AOSLO raster. The subject moved a 3-arcmin gray square on the monitor until it was centered in a 5-arcmin square rendered at the center of the AOSLO raster. B. Eye movements during the free-viewing task overlaid on the image that the subject viewed. The subject exhibited microsaccades and drifts while exploring the image during a 30-second viewing interval. C. Eye movements during the high-acuity task overlaid on the eye chart image that the subject viewed. Each line of the eyechart was viewed during a 10-second interval D. Time-course of the eye movements during the high-acuity task trimmed to 4-seconds for each line. The three rows correspond to the three rows of the eye chart, and the y-axis scale is adjusted for each line to account for the difference in microsaccade amplitude between the rows. The vertical magenta bars denote microsaccades.

To evaluate the precision of this procedure, we conducted three or more repeated measurements at each of six different time points over the course of the 15-minute eye-tracking experiment, for a total of 19 measurements of the monitor pixel coordinates corresponding to the center of the AOSLO raster scan. The standard deviation of the 19 repeated measurements was 0.41 arcmin for horizontal and 0.65 arcmin for vertical, and there was no systematic drift of the center coordinates over time. These results demonstrate alignment between the AOSLO raster and the monitor with sub-arcmin uncertainty.

In the free-viewing task, the subject freely explored images of natural scenes and city landscapes. Each image was presented for 30 seconds and subtended 2 degrees by 1.5 degrees of visual angle. The imaging wavelength of the AOSLO was 840 nm and the field of view was 1.5 x 1.5 degrees. An example of the subject’s eye movements recorded in this task is shown in Fig. 8B.

In the high-acuity task, the subject examined three lines of a tumbling E eye chart. Each of the three lines of the eye chart contained six E optotypes with randomly distributed orientations. The whole eye chart image spanned 1.47 x 0.74 degrees of visual angle and was presented at full contrast on a white background on the monitor. Stroke widths ranged from 1.1 arcmin to 0.6 arcmin across the three lines, with center-to-center spacings between 16.6 arcmin and 8.3 arcmin. The subject viewed each line of the eye chart twice, and a 10-second video of the retina was recorded during each viewing period. Fig. 8C shows the results from the high-acuity task, with eye movements from each of the three lines superimposed on the eye chart image. Eye movement traces are shown in Fig. 8D for 4-second excerpts from the full eye movement data collected on each line of the eye chart.

The recorded retinal videos were analyzed offline using a registration and motion extraction algorithm developed by Stevenson, Roorda, and Kumar [23]. The extracted eyetraces, which were produced during the video registration process, were then aligned to the coordinate system defined by the center of the AOSLO raster by independently tracking the motion of the preferred retinal locus of fixation (PRL) in the raw videos using normalized cross-correlation at the video frame rate of 30 Hz. The PRL and corresponding high-resolution retinal image for this subject was previously acquired during a separate AOSLO psychophysics experiment. The measured uncertainty during the eyetrace alignment procedure was 0.19 arcmin for the horizontal direction and 0.14 arcmin for the vertical direction. This uncertainty was quantified as the standard deviation of the difference in pixel coordinates between the extracted PRL position and the eyetrace pixel coordinates. Each pixel subtends 0.18 arcmin in the AOSLO raster, so the measured alignment uncertainty is comparable to the pixel size in both the horizontal and vertical directions.

The results shown in Fig. 8 highlight the precise nature of fixational eye movements. Consistent with results from Intoy and Rucci [52], this subject exhibited precise microsaccades with amplitudes smaller than 10 arcmin for the smallest optotypes. In between microsaccades, periods of ocular drift moved the stimulus over many photoreceptors. By simultaneously imaging the retina with cellular resolution during these eye-tracking experiments, it is now possible to track cone locations during the task, opening the possibility to generate cone activation maps in both space and time in future studies.

## 6. Discussion

We present a novel combination of a high refresh rate display with an AOSLO for natural monocular viewing of complex color stimuli in AOSLO psychophysics experiments. This combined system utilizes the spatial accuracy advantages of AOSLO-based eye-tracking while greatly expanding the range of stimuli that can be presented. For example, a video recording of a natural scene could be displayed on the monitor at 360 frames per second in color while the AOSLO recorded a video of the retina, enabling high-resolution retinal imaging and eye-tracking during natural viewing tasks. This type of stimulus delivery is not possible using the AOSLO raster.

With the current hardware, most AOSLOs designed for imaging the central foveal with cellular resolution are limited to 1-2 degrees for the full field of view, with this limitation being set by the need for sufficient pixel density to resolve the smallest foveal cones. To image a larger portion of the retina, images must be captured at different eccentricities and then stitched together. Without the use of an external display for rendering fixation markers, experimenters can instruct participants to fixate on the corners of the raster. This strategy of using the raster corners as fixation markers increases the effective field of view by a factor of approximately two and increases the retinal area of the image by approximately four times the area of the AOSLO raster [10,53]. By using an external fixation marker and automatic montaging of retinal images, it is possible to construct a composite retinal image that covers a substantially larger area of the retina while maintaining high resolution.

Automatic retinal image montaging has been demonstrated in prior art, but most implementations have focused on obtaining image strips that extend into the periphery along two cardinal axes rather than constructing continuous retinal montages that encompass the entire 5-degree fovea [49,54,55]. For quantifying cone density across the entire fovea, it is advantageous to obtain a continuous map of the photoreceptor mosaic with a diameter larger than 5 degrees centered on the foveola, which is why we implemented the full-field imaging strategy described in Section 5.1. Our approach yields a continuous map of the photoreceptor mosaic that covers an area more than 45 times larger than the standard AOSLO imaging field of view of 1 degree squared while meeting or exceeding the field of view, resolution, and stitching fidelity compared with other published full-field retinal montages [56,57].

A limitation of the eye-tracking approach presented in this paper is the small trackable range of 1.5 degrees, or +/-0.75 degrees from the center of the AOSLO raster, which we demonstrated in the eye-tracking results presented in Fig. 8. This limitation is not fundamental, however, and the trackable range could be increased by modifying the offline video registration algorithms. By using a large external reference image—such as the montage shown in Fig. 7—the trackable range could potentially be expanded to cover the whole field of view of the monitor. High-resolution Dual-Purkinje Image (DPI) eye-trackers achieve trackable ranges between +/-10 degrees and +/-25 degrees with arcminute uncertainty [58,59], but these eye-trackers rely on external calibrations to determine the position of the line of sight in the world rather than tracking retinal movement directly. TSLOs can achieve a trackable range around +/-3 degrees by trading off spatial resolution with field of view, but these systems lack the resolution for simultaneous cone imaging in the central fovea [24]. The primary application of this combined display and retinal imaging system is for studying fixational eye movements at high resolution, and thus the current trackable range of +/-0.75 degrees is sufficient for measuring both drifts and microsaccades—small, rapid gaze shifts with amplitudes less than 30 arcmin [60]— with high resolution. Future work with the video registration and eye-tracking algorithms is poised to yield an increased trackable range.

By continuously monitoring the retinal location of a projected stimulus, AOSLO- and TSLO-based eye-trackers can directly measure the retinal motion of a stimulus. This direct measurement provides knowledge of where the stimulus falls on the retina that is not available with other eye-tracking methods such as DPI and pupil tracking. The combined display and AOSLO system that we describe in this paper capitalizes on this feature of AOSLO-based eye-tracking while enabling flexible visual stimulation by means of an external monitor.

By moving the stimulus display out of the AOSLO raster and onto an external display, an additional alignment uncertainty is introduced, which is the registration between the AOSLO raster and the monitor pixels. The calibration procedure for registering the two coordinate systems, described in Section 5.2, ensures that the alignment uncertainty is maintained below 1 arcmin. Over the course of the eye-tracking experiments that we conducted, the measured alignment uncertainty between the AOSLO and the monitor was +/-0.77 arcmin, which accounts for the variance in both the horizontal and vertical directions. The other source of uncertainty in measuring the gaze position is the eyetrace extraction, which we measured to be +/-0.24 arcmin—accounting for both horizontal and vertical variance—by comparing the extracted eyetrace data with an independently tracked retinal location in the raw videos, as described in Section 5.2. After adding the variances for these two alignment uncertainties, we find that the total alignment uncertainty is +/-0.80 arcmin. This finding demonstrates that we achieve eye-tracking with sub-arcminute uncertainty using an external display for stimulus presentation and an AOSLO for eye-tracking.

With the flexible stimulus delivery capabilities of this combined system, we provide advanced functionality for investigating the spatial and temporal properties of human vision, with applications in both basic science and clinical work. Some applications include assessing visual acuity under normal viewing conditions at different eccentricities to explore the relationship between cone density, retinal image motion, and acuity; studying the temporal sensitivity of the human retina at different eccentricities; and identifying the preferred retinal locus of fixation in more natural viewing conditions and in relation to microsaccade behavior. In future studies, this external monitor can be coupled with gaze-contingent display technology for retinal stabilization and other gaze-contingent human psychophysics experiments [52,60,61] to further expand the capabilities of AOSLO-based eye-tracking.

## 7. Conclusion

We present the integration of a high refresh rate display with an AOSLO, which enables color stimulus delivery at high spatial and temporal resolution in AOSLO psychophysics experiments. Details of the design, assembly, and validation of the display are highlighted. We demonstrate high-resolution human retinal imaging over the full field of view of the display for one individual with low myopia (i.e., -1 diopter) and normal vision and construct a continuous full-field retinal montage that encompasses the entire fovea. Two eye-tracking experiments were conducted to demonstrate the advanced display capabilities that go beyond what is possible using the AOSLO raster to render stimuli. We tracked eye movements with sub-arcminute uncertainty during a free-viewing task of a color image and during a high-acuity eye chart task.

## Acknowledgements

We thank Krishnamachari S. Prahalad and Paul C. Jolly for helpful comments and discussions regarding custom software that we used for the eye-tracking experiments. We thank the technical staff at the Center for Visual Science at the University of Rochester for fabricating custom mechanical components necessary for assembling the custom optical relay. We thank Synopsys, Inc. for student licenses of CODE V, and we thank Onshape for student licenses for CAD modeling software.

## Data availability

Data files underlying the results in this paper are available upon reasonable request.

